# Untargeted metabolite analysis of *Ocimum* leaves shows species specific variations

**DOI:** 10.1101/673269

**Authors:** Manu Shree, Ranjan K Nanda, Shyam K Masakapalli

**Affiliations:** School of Basic Sciences, Indian Institute of Technology, Mandi, Himachal Pradesh, India. 175005; Translational Health Group, International centre for Genetic Engineering and Biotechnology (ICGEB), New Delhi, India, 110067

**Author notes:** Corresponding authors Shyam Kumar Masakapalli (PhD), School of Basic Sciences, Indian Institute of Technology, Mandi, Himachal Pradesh, India. 175005., E. mail or, Ranjan Kumar Nanda (PhD), Translational Health Group, International center for Genetic Engineering and Biotechnology (ICGEB), New Delhi, India. 110067., E. mail.

**Keywords:** *Ocimum*, *O. basilicum*, *O. kilimandscharicum*, *O. sanctum*, metabolite profiling, metabolomics, GC-MS

## Abstract

Tulsi (*Ocimum* species), the queen of herbs is a common ingredient in beverages with perceived health benefits. Recently published *Ocimum* genome highlighted the presence of several genes that contributes to important phytochemicals but a comprehensive metabolite profiling to study the water soluble metabolites of *Ocimum* is lacking. In this study, untargeted metabolic profiling of hot water extract of fresh and air dried leaves of *O. basilicum, O. sanctum* and *O. kilimandscharicum* species employing gas chromatography and mass spectrometry (GC-MS) was attempted. Analysis of hot water extracts of *Ocimum* leaves will provide details of molecules consumed and species specific differences, if any. Several metabolic features including amino acids (glycine, serine glutamate), organic and other acids (succinic acid, fumaric acid, 4-amino butanoic acid), sugars and their derivatives (glucose, sucrose, mannitol, fructose) and secondary metabolites (shikimic acid, quinic acid, catechol, gamma amino butyric acid, eugenol) were identified. Multivariate statistical analysis of GC-MS data indicated several species specific metabolic similarities and differences. Based on variable importance parameter score of >1, it was observed that in case of air dried extracts, glucose, fumaric acid, and D-mannitol displayed as important variables for species specific variation. Whereas in case of fresh leaves extracts, the variation was prominent due to xylose, D-allose and an unknown metabolic feature detected at 24 min (metabolite@24 with highest m/z 75). Phytochemical phenotype of *Ocimum* leaves not only shows species specific variations but these may partly explain their difference in taste and health benefits from their use as hot beverages.

## INTRODUCTION

Plants are rich source of phytochemicals with promising health and industrial applications. Plant parts like fresh or dried leaves, stems and roots are commonly used by humans for beverage prepration with perceived health benefits. Profiling micromolecules of these plant materials could be of immense interest to generate evidence to explain the health benefits and may also be useful for strain improvement. *Ocimum* genus belongs to lamiaceae family, comprises of more than 200 species of herbs and shrubs with immense medicinal values^1,2,3,4^. It is widely used for preparing beverage in South East Asia including India. The phytochemical studies on *Ocimum* species revealed that these are rich source of terpenoids and phenolic compounds including phenolic acids, phenols, and polyphenols such as flavonoids and anthocyanins^5^. In addition, recently published genome sequence of tulsi highlighted the presence of genes encoding pathways related to several health beneficial phytochemicals such as apigenin, ursolic acid and taxol^6^. Very limited reports available in literature explaining the metabolites of hot water fresh leaf and dried leaf extracts from different *Ocimum* species.

Metabolomics deal with studying the complete set of metabolites (small molecules) present in a cellular system. Measuring metabolite profiles at different conditions capture variations or similarities in standard and perturbed physiological conditions. These profiles are primarily concerned with the global or system-wide characterization of all metabolites present in the samples using technologies such as nuclear magnetic resonance (NMR), liquid/gas chromatography and mass spectrometry (LC/GC-MS). Common analytical platforms with better sensitivity like Gas Chromatography-Mass Spectroscopy (GC-MS) is widely used for metabolite profiling from diverse biological samples including plant material. The process involves derivatization of the phytochemicals along with internal standards (like ribitol, lysine-D4 etc) to volatilize and detect by GC-MS. Methoximation and Silylation derivatization method is routinely used to cover wide range of metabolites including amino acids, alcohols, sugars, amines, sugar alcohols, secondary metabolites ^7^.

In this study, we aimed to carry out global metabolite profiling to study metabolites extracted from fresh and air-dried leaves of three *Ocimum species (O. basilicum, O. sanctum* and *O. kilimandscharicum)* using GC-MS (Figure 1). Dried or fresh leaves of several *Ocimum* species are used as ingredients in beverages (as Tulsi tea), ayurvedic formulations and culinary practice. Evaluating the metabolic variations and similarities between the hot water extracts of air-dried and fresh leaves and also among the *Ocimum* species would be of interest to understand the repertoire of health relevant phytochemicals that are being consumed by the end-users.

**Figure1:**
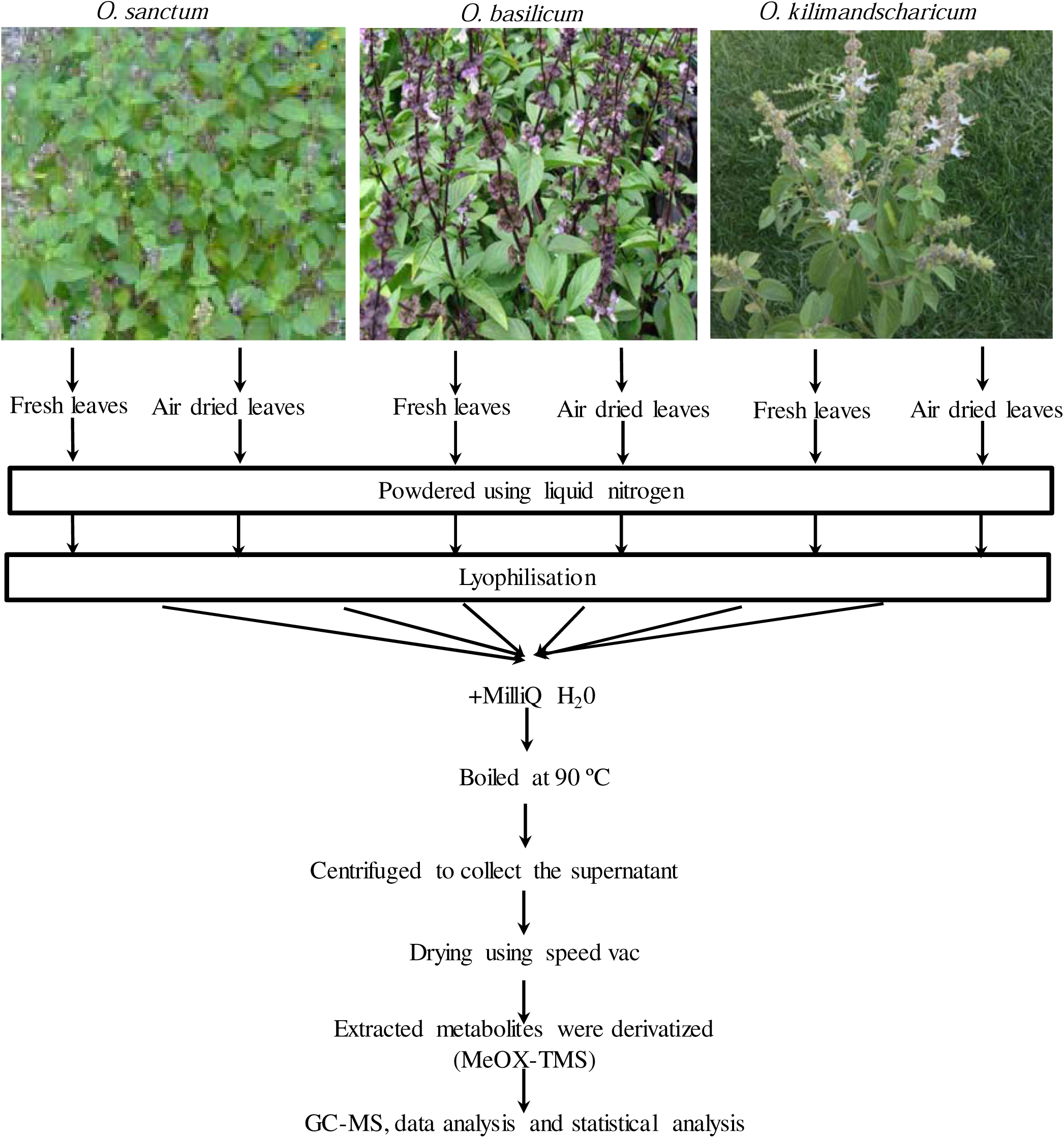
Experimental work flow. Leaf samples of field grown *Ocimum* species (*O. sanctum, O. basilicum, O. kilimandscharicum*) were harvested followed by further processing for metabolite profiling. Air dried leaves were obtained by shade drying. The fresh leaves harvested were immediately quenched in the field using liquid nitrogen. The samples stored in −80 °C were subjected to hot water extract and subsequent derivatisation for GC-MS

## MATERIALS AND METHODS

### Plant materials

*O. sanctum, O. basilicum and O. kilimandscharicum* plants grown in the medicinal plant garden of Indian Institute Technology Mandi, Kamand located in the mid-Himalayan region (altitude: 972 meters mean sea level) were washed sprinkling water. After carefully washing the handpicked leaves were placed in zipped lock poly bags in five per set. Each set was either used fresh or kept for air-drying. A set of fresh leaves were dipped in liquid nitrogen and powdered using mortar and pestle before lyophilization. The other set was air-dried under shade for 15 days at room temperature and liquid nitrogen was used for powdering before lyophilization. The lyophilized samples were stored in air-tight containers in −80 ºC until further analysis as outlined in Figure 1.

### Gas Chromatography Mass Spectroscopy(GC-MS)

The hot water infusion extracts (n=5) were obtained by adding 1 ml of H_2_O to 50 mg of air-dried and fresh samples followed by heating at 90 ºC for 10 min. The extracts were subjected to centrifugation at 13,000 × g for 10 min at 4ºC to separate the water-soluble metabolites from insoluble pellets. The supernatant containing water soluble metabolites was then mixed with ribitol (10 μg/μl), as an internal standard to access the quality of derivatisation and metabolite quantification. The extracted polar metabolites (100 μl) and internal standard mix was then collected into fresh tube and was dried by vacuum-drier at 40 ºC for 2 hours using a speedvac (Labconco, USA). The dried extracts were subjected to methoxyamine hydrochloride (MeOX)- Trimethylsilyl (TMS) derivatization^8^. Briefly, to each dried extract 35μl of methoxyamine hydrochloride (20 mg/ml in Pyridine) was added and incubated for 2 h at 37 ºC and 900 rpm on a thermomixer. After adding 49ul of MSTFA (mono-trimethylsilyl-trifluoroacetamide) to each sample mixture were incubated at 37 ºC for 30 min at 900 rpm in a thermomixer. The supernatant, after centrifugation at 13,000 × g for 10 min were transferred to a glass vial insert for GC-MS data acquisition. The derivatised samples from each treatment condition (fresh and air-dried) were analyzed in the same run to maintain consistent parameters for all the samples throughout the procedure. The metabolite stability, chromatographic peaks and their reproducibility was ensured by the use of internal standard ribitol in a concentration of 0.01 μg/μl in each of the derivatised samples. Occurrence of chromatographic peak for ribitol at a retention time of 21 min in all the samples ensured consistency of the runs. The instrument detection limit (IDL) has been tested by injecting with octafluoronaphthalene (100 fg/ μl) and the IDL limit was found to be 2 fg. For confirming identity of molecules, commercial standards of myoinositol, glycine, serine, shikimic acid, fumaric acid, tartaric acid, quinic acid, and 4-amino butyric acid (Sigma-aldrich, USA) were subjected to derivatization following similar methods used for sample processing before GC-MS analysis.

### GC-MS Data acquisition

Derivatized samples (1μl) were injected, using an automatic liquid sampler (7683B ALS, Agilent, USA) by splitless mode into GC-MS system (7890A GC, 5975, USA). Molecules were separated by an DB-5MS column (5% diphenyl, 95% dimethylpolysiloxane; 30 m × 0.25 mm × 0.25 μm; Restek USA) and helium was used as a carrier gas at a constant flow rate of 1 mL/min. Front injection temperature was set constant at 250 °C. A temperature gradient of 1 min at 50 °C isothermal, followed by a 10 °C/min oven temperature gradient to a final 200 °C, and then hold for 4 min at 200 °C, followed by a 10 °C/min to a final 300 °C temperature for 10 min was used. Ions were generated by a 70 eV electron beam in electron ionization mode. The spectra were recorded with a scanning range of 50-600 mz^-1^. The MassHunter software (Agilent Technologies, USA) was used to control the data acquisition parameters (both GC separation and mass spectrometry) during sample introduction and the total run time was 50 min per sample. The lower limit of detection of the adopted method was calculated to 0.01 µg/µl of ribitol. All analysis of the raw data files is presented in files S1 and S2.

### Data pre-treatment

In order to reduce noise in spectra due to GC-MS instrument, the raw spectra were subjected to baseline correction using Metalign software^9^. Peaks of baselined spectra were identified using MassHunter software based on MS search from NIST17 and Fiehn13 library (NIST, National Institute of Standards and Technology, Maryland, Golm database). The peak integrals were obtained using the MassHunter Qualitative Navigator B.08.00 software (Agilent, USA)^10^. The peaks between spectra were compared based on the m/z of different fragments, their elution times and hits against the library using MassHunter Qualitative Workflows B.08.00 software (Agilent, USA). The metadata file compiled from the GC-MS spectral analysis of the three *Ocimum* species including the metabolite identified, elution time and m/z fragmentation patterns was used for multivariate statistical analysis. All metadata files generated from fresh and air-dried *Ocimum* samples are tabulated in files S1 and S2 respectively. The relative fold changes of the identified metabolites were calculated with respect to the internal standard Ribitol.

### Statistical analysis

MetaboAnalyst 4.0 was used for multivariate statistical analysis using GC-MS meta data^11^. The Log transformation and pareto scaling method of data normalization were selected for analysis ^12,13^. Principal component analysis (PCA) was carried out to assess the differences in samples preparation and heterogeneity between different *Ocimum* species^14^. In addition, molecules with variable importance in projection (VIP) score of >1.0 were identified as important metabolites for analysis of species-specific variation. In order to determine the significant differences between and within the *Ocimum* varieties based on the identified metabolites under air-dried and fresh treatment conditions, one-way analysis of variance (ANOVA) and post hoc analysis using Tukey’s Honestly Significant differences (Tukey’s HSD) and False discovery rate (FDR) was performed.

## RESULTS

### Metabolite profiles of *Ocimum* species

The total ion chromatograms (TIC) of fresh and air-dried *Ocimum* leaves hot water extracts generated by GC-MS (figure 2a). A total of 144 GC-MS peaks of MeOX-TMS derivatised extracts (fresh and air-dried) were annotated (Table S1 and S2) as metabolic features. The metabolic features with a probability score of >70% against the NIST17 and Fiehn13 library and/or those that matched the elution times and m/z fragmentation patterns of the metabolic standards were considered identified. Among these, 14 peaks were unknown and labelled as “metabolite @retention time”. The identified metabolites (n=130) from all the extract were grouped into organic acids (malic, fumaric, lactic, and tartaric acid), sugars and their derivatives (glucose, xylose, fructose and mannitol) and secondary metabolites and their pathway intermediates (shikimic acid, quinic acid, myoinositol and eugenol).

**Figure 2:**
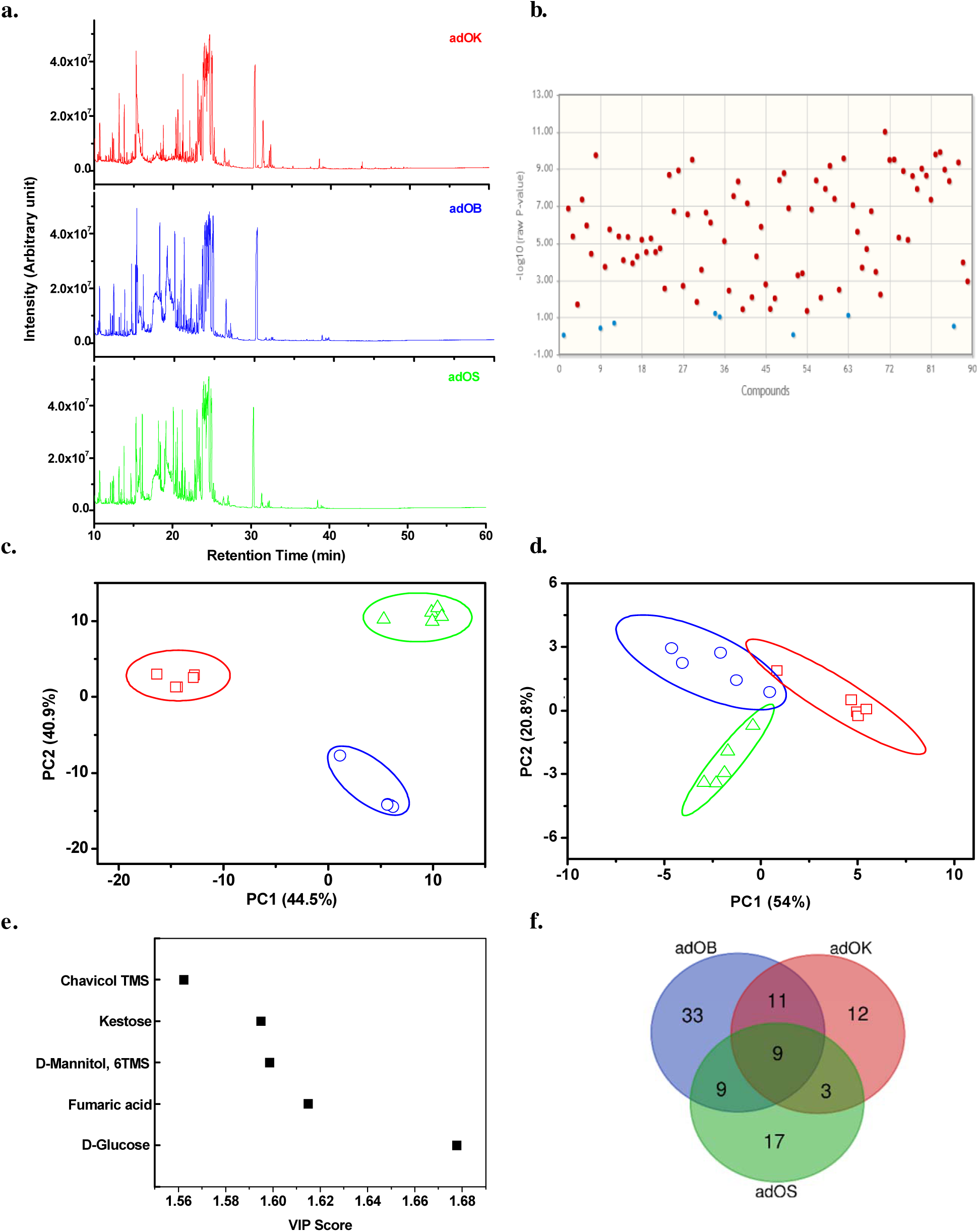
Metabolite profiling and metabolomics of hot-water extracts of air dried leaves of *Ocimum species*. The Total ion chromatograms of air-dried leaves of *O. basilicum* (adOB); *O. kilimandscharicum* (adOK); *O. sanctum* (adOS). (b) One way ANOVA showing significant metabolites with p value < 0.05 among all the air-dried samples (c) Principal component analysis (PCA) plot of all the metabolic features and (d) only the common metabolic1 features (e) Variable importance in projection (VIP) score plot showing the main metabolic features that contribute towards variance among the *Ocimum* species. (f) Venn diagram depicting the distribution of metabolic features between and within the *Ocimum* species.

### Metabolic variations and similarities in air-dried leaves of *Ocimum* species

The GC-MS based metabolite profiles of the air-dried (Figure 2a) samples of *O. basilicum, O. sanctum and O. kilimandscharicum* shared differences and similarities among the species. A total of 86 metabolic features out of 95 were found to significantly vary among the three *Ocimum* species as confirmed by one-way analysis of variance (ANOVA) and Tukey’s Honestly Significant Difference (Tukey’s HSD) based post-hoc tests (Figure 2b and Table S3a). Unsupervised multivariate data analysis of all the metabolic features from air-dried leaf samples by PCA resulted in different clusters of *O. basilicum, O. sanctum and O. kilimandscharicum* with PC1 and PC2 scores of 44.5% and 40.9% respectively (Figure 2c). Common metabolites such as succinic acid, 4-amino butanoic acid, quinic acid, fructose, malic acid and other unknowns were detected in the air-dried extracts across *Ocimum* species. PCA analysis of 9 common metabolic features (PC1 and PC2 contributed to 54.1 and 20.8% variance respectively), clustered the *O.basilicum* species separately (Figure 2d). Among the common metabolic features in all the species, only 4 were found to significantly vary as revealed by ANOVA (p≤0.05, FDR≤0.1) and Tukey’s Honestly Significant Difference (Tukey’s HSD) based post-hoc tests (Table S3b). The metabolic features contributing towards species specific variations based on VIP scores (Figure 2e) are predominantly D-glucose, Fumaric acid, D-mannitol etc. A Venn diagram showing distribution of the number of common and unique metabolic features between and within the *Ocimum* species is provided in figure 2f and Table S3c.

### Metabolic variations and similarities in fresh leaf samples of *Ocimum* species

The metabolite profiles of fresh *Ocimum* samples were obtained using GC-MS as shown in figure 3a. In case of fresh leaf extracts, a total of 91 out of a total of 106 metabolic features significantly altered among all the three *Ocimum* species as revealed by ANOVA and Tukey’s Honestly Significant Difference (Tukey’s HSD) based post-hoc tests (Figure 3b and Table S4a). Unsupervised multivariate PCA of all identified metabolic features (PC1 48.4% and PC2 31.7%) captured distinct variations among the *Ocimum* species (Figure 3c). A total of 8 common metabolic features were present in fresh leaf extracts including shikimate, fumarate, quinic acid, tartaric acid, myoinositol and others. These common metabolic features also subjected to PCA (PC1 50.1% and PC2 23.2%) showed clear distinction among the *Ocimum* species (Figure 3d). Among these, 3 metabolites levels significantly different among the *Ocimum* species as confirmed by one-way ANOVA (p≤0.05, FDR≤0.1) (Table S4b). Among all the metabolic features analysed, Xylose, D-allose, D-fucitol are found as important molecules (VIP > 1.0) that explain the variations among *Ocimum* species (Figure 3e). A Venn diagram showing the distribution of the number of common and unique metabolic features between and within the *Ocimum* species is presented in Figure 3f and Table S4c.

**Figure 3:**
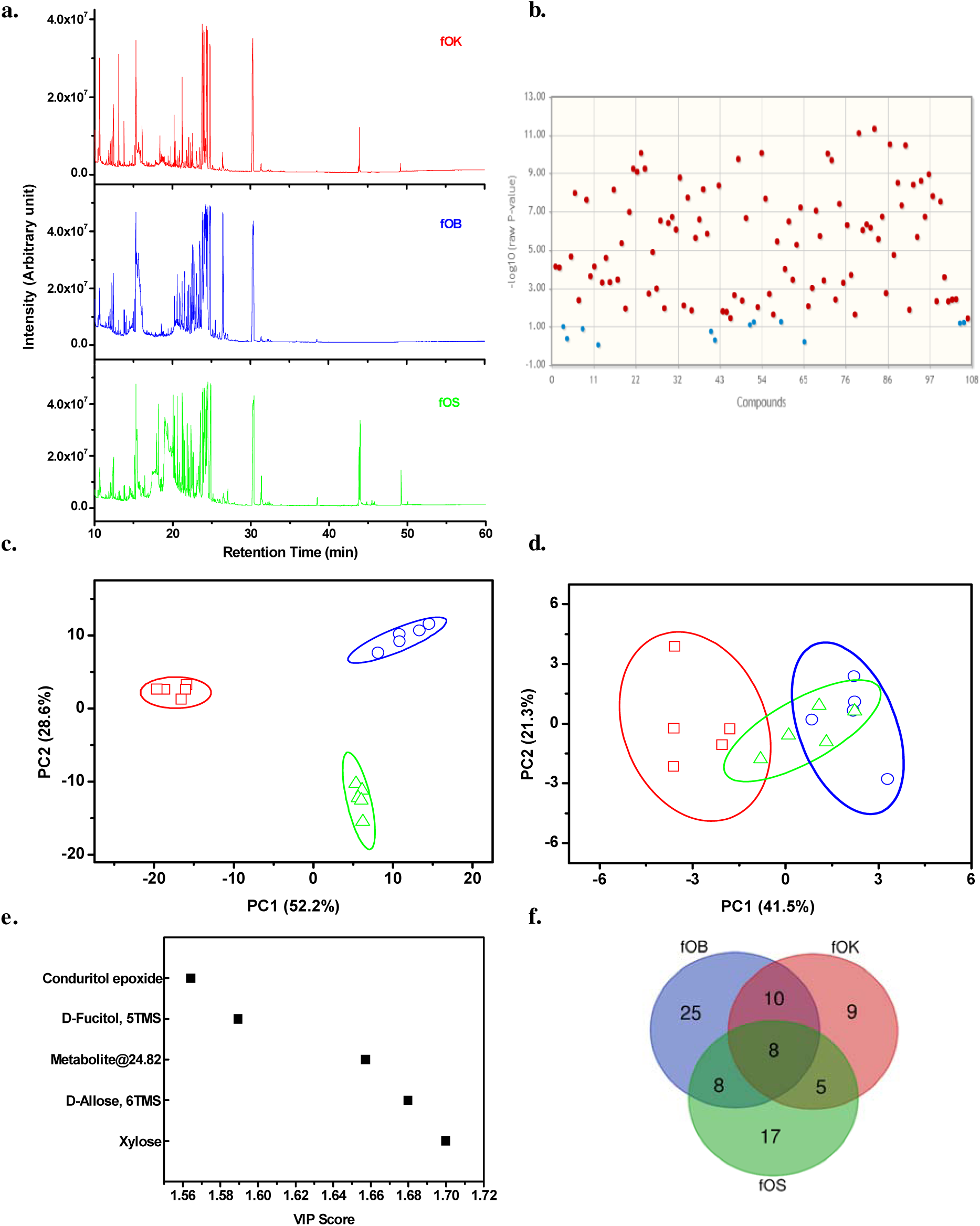
Metabolite profiling and metabolomics of hot-water extracts of fresh leaves of*Ocimum species*. (a) The Total ion chromatograms of air-dried leaves of *O. basilicum* (fOB); *O. kilimandscharicum* (fOK) *O. sanctum* (fOS). (b) One-way ANOVA showing significant metabolites with p value < 0.05 among all the fresh samples. c) Principal component analysis (PCA) plot of all the metabolic features and (d) only the common metabolic features. e) Variable importance in projection (VIP) score plot showing the main metabolic features that contribute towards variance among the *Ocimum* species. (f) Venn diagram depicting the distribution of metabolic features between and within the *Ocimum* 1 species.

### Metabolic network of identified molecules

All identified metabolites from the hot water extracts of *Ocimum* species (*O. basilicum, O. sanctum* and *O. kilimandscharicum*) were overlaid to report the active central and secondary metabolic pathways in *Ocimum* leaves (Figure 4). The precursor, intermediates and product metabolites detected in *Ocimum* leaf extracts have could be mapped to central (Glycolysis, Pentose phosphate pathway, TCA cycle, GABA shunt) and secondary metabolic pathways (Shikimate Pathway, Phenyl propanoid pathway and others). Secondary metabolites like shikimic acid, quinic acid, myoinositol, catechol, GABA: gamma amino butyric acid, eugenol and tartaric acid showed species specific variations (Figure 1). Shikimic acid abundance measured in the three *Ocimum* varieties under fresh and air-dried conditions were found to be statistically different (ANOVA, p value < 0.05). Several other health beneficial metabolites showed significant variation among different *Ocimum* varieties. Myoinositol, tartaric acid (p value < 0.005) and catechol (p value < 0.05) was detected in case of air-dried *O. sanctum* and *O. basilicum*. Whereas quinic acid (p value < 0.05) was found to be significantly present in both airdried *O. sanctum* and O. *basilicum.* In case of fresh *Ocimum* leaf extracts, eugenol was only detected in *Ocimum sanctum* (p value < 0.0005) and GABA in only *O. basilicum* (p value < 0.005). Fresh leaf extracts from *O. basilicum* and *O. kilimandscharicum* also contained significant levels of catechol (p value < 0.05) and quinic acid (p value < 0.005) as shown in figure S1.

**Figure 4:**
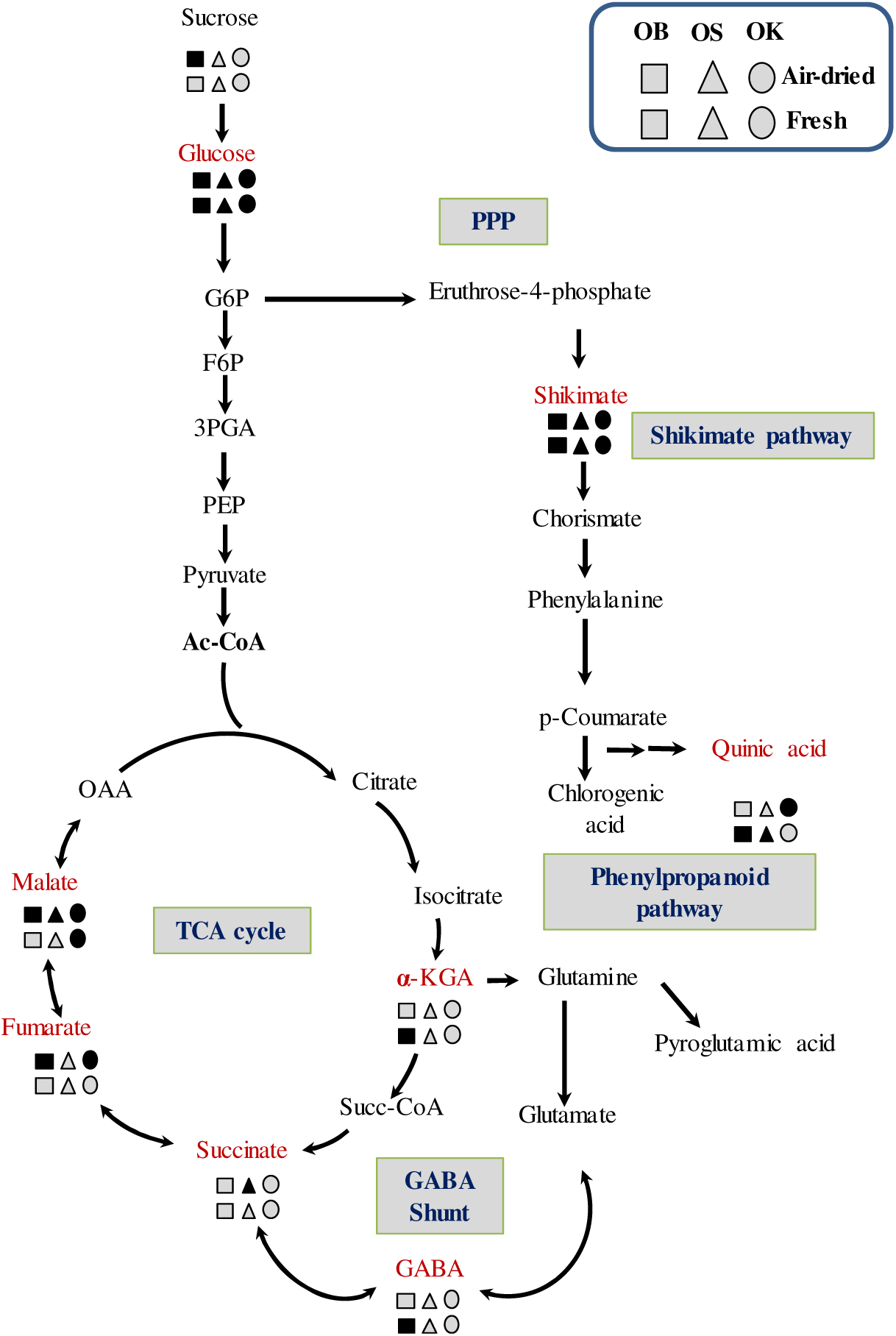
Mapping of identified metabolites of *Ocimum* on to a simplified central and secondary metabolic network. The identified metabolites of the three *Ocimum* varieties are highlighted in red. Squares, triangle and circles represent *O. basilicum, O. sanctum* and *O. kilimandscharicum* metabolites respectively from air dried (top row) and fresh (bottom row) leaf extracts. Black and grey shades in the symbols represent the metabolite as detected or undetected in the hot water extracts as profiled from the GC-MS analysis.

## Discussion

*Ocimum*, the queen of herbs, being consumed since ages comprises multiple health beneficial metabolites. Earlier studies extracted metabolites from *Ocimum* using organic solvents like methanol, ethanol and demonstrated their medicinal values^15,16,17,18,19^. These molecules (∼30) were from different classes of flavonoids, sesquiterpenes, terpenes, phenylpropanoids and sterols ^6^. Molecules like eugenol, ursolic acid, apigenin and rosmarinic acid are well characterized on their medicinal property and are extracted in ethanol/methanol. In addition, these molecules also show Ocimum species specific abundance and demonstrate solvent specific recovery. With several of these key metabolites preferential solubility to organic solvent, it is of interest to investigate the metabolic composition of the users consuming hot aqueous beverages made from *Ocimum* leaves. These molecular details are hard to find in literature and profiling these molecules, may be useful to understand the molecules responsible for the taste and perceived health benefits of hot Tulsi (*Ocimum*) leaf tea, a widely accepted beverage. In addition, the hot water beverages from different *Ocimum* species are anticipated to have their different taste and may partly be contributed by varied method of processing and in the metabolite profiles.

In the current study, GC-MS based global metabolites of hot water leaf extracts of *O. basilicum, O. sanctum* and *O. kilimandscharicum* was employed. Several metabolic features including amino acids (glycine, serine glutamate), organic and other acids (malic acid, succinic acid, fumaric acid, 4-amino butanoic acid), sugars and their derivatives (glucose, galactinol) and unclassified secondary metabolites and their pathway intermediates (ferulic acid, trans caffeic acid, shikimic acid, quinic acid, myoinositol, tartaric acid) with few unknown molecules were identified. The identities of the important metabolites were validated by running commercial standards. The dried powder and liquid concentrates of *Ocimum* leaves from different species are commercially marketed widely for their utility as infusions/teas and the current analysis augments their potential.

Our findings also corroborate earlier report on presence of malic and tartaric acid in the aqueous extract of *O*. *basilicum*^20^. In fact, these two metabolites were identified in leaf extracts of all three studied *Ocimum* varieties. In addition, *Casanova et al., 2016* also reported cinnamic acid derivatives like caftaric acid, caffeic acid, chicoric acid and rosmarinic acid in fresh leaf extracts of *O*. *basilicum*^21^. In this study, we did not identify these molecules and may be due to the formation of phenolic compounds from different chemical moieties e.g. chicoric acid is a tartaric acid ester of two caffeic acids (a hydroxy cinnamic acid) and known to be present in many plant families^22^. Chicoric and caffeic acid were not identified but tartaric acid has been reported in different hot water extracts. The presence of precursor moieties i.e. tartaric acid, explains that there might be other parameters that hindered the formation of final product i.e. chicoric acid. Sample processing and analytical techniques used for molecular detail extraction may also impact the extraction of these molecules. Moreover, temperature play a critical role in retention of phenolics in extracted plant materials. Kim et al., 2010 showed that higher air-drying temperature accelerate the loss of chicoric acid in *E. purpurea* flowers. Temperature used for preparing hot tea may be one of the probable reason of not identifying caftaric acid, chicoric acid and rosmarinic acid in our analysis^23^. Drying temperature of plant materials could play a role in retention of phenolics in plant materials. The air-drying temperature (70 °C compared to 40 °C and 25 °C) may be caused loss of phenolic compounds in our study^23^. In order to achieve complete validation for their absence, MeOX-TMS derivatised commercially available standards were run in GC-MS. We did not identify any of these molecules like caftaric acid, caffeic acid, chicoric acid, rosmarinic acid and ursolic acid in our analysis so highly likely we are losing them during sample processing (data shown in table S1 & S2).

To the best of our knowledge, not much literature information is available on metabolic constituents of hot water extracts of *O. sanctum* and *O. kilimandscharicum* and the current study provides insights into it for the first time. These species showed similar and unique metabolites in two different treatment conditions i.e. the air-dried and fresh *Ocimum* leaves. GC-MS based metabolite profiles highlighted the presence of health beneficial secondary metabolites (shikimic, catechol, eugenol, GABA etc.) more prominently in fresh leaves as compared to the air-dried ones and hence could be of interest to the consumers^24,25^. It is important to mention that fresh leaves may be yielding better nutriprotective molecules and could be a better target for use. Several of the identified metabolites (such as shikimic acid, tartaric acid, myoinositol, quinic acid, GABA etc.) can be correlated to various health benefits like antioxidant, anti-diabetic and immunomodulatory properties as explained in scientific literature^19,26^. Our study confirmed the presence of different health beneficial metabolites in all the three *Ocimum* leaf varieties.

## Conclusion

In summary, the metabolite profiles of the hot water extracts from three *Ocimum* species (*O. basilicum, O. sanctum, O. kilimandscharicum*) are documented using GC-MS. The molecular similarities and differences among the *Ocimum* species and the processing conditions of air-dried and fresh leaves were analyzed. In particular, species specific variation was observed and statistically validated based on the metabolite profiles of *O. basilicum, O. kilimandscharicum* and *O. sanctum.* The metabolite profiles of the *Ocimum* species can be correlated to various health benefits they offer when consumed as hot water beverage and few of these molecules could be useful as marker for quality control checks for finding batch to batch variation in industrial production.

## Author information Contributions

S.K.M and M.S. designed the study. M.S. performed the experiments. M.S., R.K.N. and S.K.M analysed the data and wrote the manuscript.

## Competing interests

The authors declare no competing financial interests.

## Acknowledgements

SKM acknowledges funding from IIT Mandi (IITM/SG/SKM/48) and SERB (ECR/2016/001176). Also BioX center and AMRC, IIT Mandi are acknowledged for access to facilities.

