## Supplementary material for "Untargeted metabolite analysis of *Ocimum* leaves shows species specific variations": Figure S1

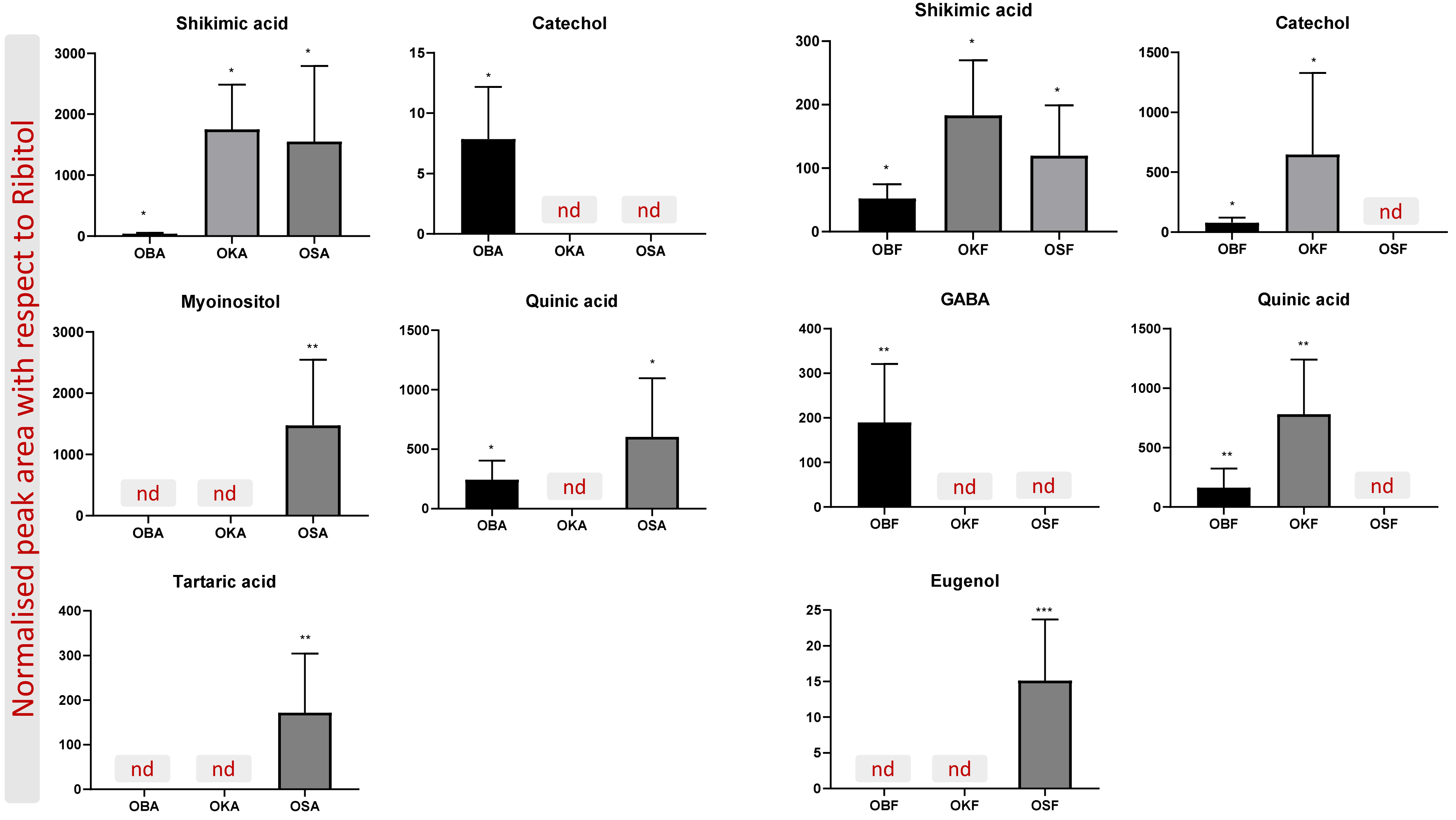


**Figure S1:** Relative fold changes and the detection limits of some health beneficial metabolites with respect to the internal standard ribitol under air-dried (Shikimic acid, catechol, myoinositol, quinic acid, tartaric acid) and fresh(shikimic acid, catechol, GABA: gamma amino butyric acid, quinic acid and eugenol) treatment condition. The air-dried *Ocimum* samples are represented as: OBA - air dried *Ocimum basilicum* ; OKA- air dried *Ocimum kilimandscharicum*; OSA - air dried *Ocimum sanctum* and fresh *Ocimum* samples are shown as: OBF – fresh *Ocimum basilicum* ; OKF- fresh *Ocimum kilimandscharicum*; OSF - fresh *Ocimum sanctum.* Asterisks represent statistically significant differences (***, $P < 0.0005$; **, $P< 0.005$ and *, $P < 0.05$) as determined by one-way ANOVA; nd, undetermined. Error bar represents standard deviation (sd) of five replicates
